# Dietary nitrate does not alter the rate of torque development during maximal voluntary isometric or electrically evoked tetanic contractions in young men or women

**DOI:** 10.64898/2026.10.06.757070

**Authors:** Lenna K. Gottschild, Jay P. Pillai, Madison J. Fry, William S. Zoughaib, Richard L. Hoffman, Andrew R. Coggan

## Abstract

We determined the effects of NO_3_^−^ ingestion on voluntary and electrically evoked rate of torque development (RTD) in young men and women. Using double-blind, placebo-controlled crossover designs, 14 men or 12 women (tested during both early and late follicular phases) acutely ingested 200 μmol/kg of NO_3_^−^ from beetroot juice (BRJ) or a NO_3_^−^-free BRJ placebo. Voluntary RTD (0-30 ms, 0-50 ms, 0-200 ms, maximal) was measured during maximal voluntary isometric contractions. The maximal tetanic RTD was also determined during stimulation at 100 Hz. Muscles tested included the triceps surae and triceps brachii in men and the quadriceps femoris in women. Dietary NO_3_^−^ had no significant effect on initial or maximal voluntary RTD in men in either muscle group (P≥0.31) or in women during either menstrual phase (P≥0.69). Similarly, acute NO_3_^−^ intake did not alter maximal tetanic RTD in men (P≥0.87) or women (P≥0.70). We conclude that acute dietary NO_3_^−^ supplementation has no apparent effect on RTD during maximal voluntary or electrically evoked isometric contractions of the lower or upper limbs in young, untrained individuals. These findings argue against changes in neural activation or actomyosin crossbridge cycling rate as mechanisms driving NO_3_^−^-induced improvements in muscle contractile function.

## INTRODUCTION

Skeletal muscle contractility has historically been evaluated based on metrics such as maximal isometric strength or peak dynamic power. However, the maximal rate of torque development (RTD) has emerged as an additional important aspect of neuromuscular function (1–4). Indeed, Grossio et al. (5) recently demonstrated that such “explosiveness” independently explains nearly as much variation in muscle performance across individuals as isometric strength. Consequently, measuring RTD has become vital for providing a comprehensive assessment of the capabilities of the neuromuscular system.

Functionally, RTD is of considerable practical significance in both athletic tasks and clinical settings. In an athletic context, rapid reactive movements and postural adjustments are heavily time-constrained, with the window available to exert force frequently limited to <200 ms (2). Because maximal voluntary torque typically requires >300 ms to achieve (2), athletic performance is often fundamentally governed by RTD rather than maximal strength or power. In the clinical context, low RTD is associated with impaired physical function (6,7) and an increased risk of falls (8) in older individuals who must generate force quickly to recover from a slip, trip, or misstep (9). Interventions that can significantly increase RTD therefore have the potential to enhance sports performance, improve physical function, and minimize the risk of musculoskeletal injuries.

Mechanistically, RTD is determined by both neural and muscular factors (1–4). The early phase (<75 ms) is largely dictated by centrally mediated motor unit recruitment, as evidenced by the strong correlation between early phase RTD and integrated surface EMG activity (1). Conversely, RTD beyond this time is more intrinsically muscular, primarily reflecting the rate of actomyosin crossbridge cycling but possibly also calcium (Ca^2+^) release kinetics and myofilament Ca^2+^ sensitivity (1–4). Finally, musculotendon stiffness plays a significant role at all durations (4).

Dietary nitrate (NO□^−^) has attracted considerable attention due to its ability to enhance skeletal muscle contractile function. Ingestion of inorganic NO□^−^ results in an increase in nitric oxide (NO) production via sequential reduction of NO□^−^ to nitrite (NO□^−^) and then to NO within systemic blood and tissues (10,11). In turn, elevated NO bioavailability improves muscle speed and power (12,13), apparently by altering Ca^2+^ sensitivity in a fiber type-dependent manner (14). This could be due to phosphorylation or nitros(yl)ation) of various proteins, e.g., myosin-binding protein C, titin, etc. (15). Nonetheless, the precise mechanism(s) by which NO_3_^−^ intake modulates the contractile properties of muscle are still uncertain.

Despite potential biochemical pathways by which dietary NO_3_^−^ might influence RTD, its specific effects on this important parameter are equivocal. Haider and Folland (16) and Tillin et al. (17) initially found no effect of NO_3_^−^ supplementation on RTD during maximal voluntary isometric muscle contractions (MVICs) of the mixed fiber type knee extensors in young men. Subsequently, however, Poredoš et al. (18) reported that NO_3_^−^ intake enhanced RTD during knee extensor MVICs in young men, but not in young women. This study, however, did not control for menstrual cycle phase or hormonal contraceptive use, both of which could potentially impact the response to NO_3_^−^ supplementation (19). More recently, Wei et al. (20,21) reported that acute ingestion of a low dose of NO_3_^−^ or 2 wk of supplementation with a higher dose increased RTD during MVICs of the knee extensors in young men and women, but only at shorter durations (i.e., ≤50 ms). This is surprising, because NO_3_^−^ supplementation would not be expected to influence the neural mechanisms that primarily determine early phase RTD (1–4). Thus, whether dietary NO_3_^−^ increases RTD, and if so, how and under what conditions, is unclear.

Given the above, the purposes of the present investigation were to assess the impact of acute dietary NO□^−^ supplementation on RTD in 1) muscles differing in fiber type and 2) in women as well as men while carefully controlling for menstrual cycle phase. To differentiate between potential central neural vs. peripheral muscular mechanisms, we determined the effects of NO□^−^ ingestion on RTD during both voluntary isometric and electrically stimulated tetanic contractions.

## MATERIALS AND METHODS

### Participants and Experimental Design

This study is based on previously unreported data from two recent publications from our laboratory (14,19). Both studies used nearly identical procedures, and the complete details of these investigations are reported therein. Briefly, however, Zoughaib et al. (14) employed a randomized, double-blind, placebo-controlled crossover design, in which the effects of acute ingestion of NO_3_^−^-containing or NO_3_^−^-free beetroot juice (BRJ were determined in the triceps brachii (60-70% type II) and triceps surae (60-70% type I) muscles of healthy young men (n=14; age 26±7 y, height 1.76±0.07 m, weight 78.1±19.3 kg). Fry et al. (19) also used a randomized, double-blind, placebo-controlled crossover design, testing the effects of dietary NO_2_^−^ on the mixed quadriceps femoris muscle of healthy, normally menstruating young women (n=12; age 23±5 y, height 1.65±0.08 m, weight 64.2±13.2 kg) during both the early follicular (EF) and late follicular (LF) phases of the menstrual cycle. All participants were classified as low-to-moderately active based on the International Physical Activity Questionnaire and were screened for cardiovascular, neurological, or medication-related contraindications prior to enrollment. Participants practiced all procedures prior to their first experimental visit. Both studies were approved by the Indiana University Institutional Review Board and all participants provided written informed consent.

### Experimental Protocol

In both studies, participants reported to the laboratory following an 8 h fast and having refrained from exercising, using chewing gum or mouthwash, or from consuming foods high in NO_3_^−^ (e.g., spinach, beets, collard greens), alcohol, or caffeine for 24 h prior to the study visit. All study visits for a given participant took place at the same time of day (±1 h). An intravenous catheter was first inserted, and a blood sample was obtained for subsequent determination of plasma NO_3_^−^ and NO_2_^−^ concentrations (data reported in Refs. 14 and 19). In the female cohort, menstrual cycle phase was confirmed by subsequently also measuring plasma estradiol and progesterone concentrations (data reported in Ref. 19). The skin over the target muscles was then shaved, lightly abraded, and cleaned with alcohol to reduce impedance. Two self-adhesive electrodes (5.08 × 8.89 cm; Dura-Stick Plus, Chattanooga Medical Supply, Chattanooga, TN) were then placed over each muscle group to permit subsequent NMES as previously described (14,19). Electrode placement was individualized to minimize co-activation of surrounding musculature and documented photographically to ensure precise replication across visits. Participants then consumed 2.2 mL/kg body mass of concentrated BRJ (Beet It Sport, James White Drinks, Ipswich, UK) that either contained or was depleted of 200 µmol/kg of NO_3_^−^ and rested quietly for 2 h. Additional blood samples were obtained 1 h and 2 h following BRJ ingestion, after which muscle function testing was performed as described below, thus coinciding with peak circulating NO_3_^−^ and NO_2_^−^ levels. A final blood sample was then obtained and the participant was released.

### Muscle Function Testing

Muscle contractile function was assessed using a Biodex System 4 Pro dynamometer (Biodex Medical Systems, Shirley, NY) interfaced with a Biopac MP160 data acquisition system (Biopac, Goleta, CA) sampling at 1000 Hz (22). Participants were stabilized using waist and torso straps along with limb-specific restraints to minimize extraneous movement. The dynamometer’s axis of rotation was carefully aligned with the anatomical joint being tested. Joint configurations and positioning varied by cohort and muscle group:

1. Men (triceps brachii): Testing was performed a chair-back angle of 1.48 rad with the elbow joint fixed at 1.57 rad, the shoulder adducted, and the arm at neutral elevation (0 rad). A Roylan Ulnar Deviation Splint (Performance Health, Cedarburg, WI) and an elastic wrap were used to prevent wrist and finger movement.
2. Men (triceps surae): Testing was performed with a chair-back angle of 0.96 rad with the knee and ankle joints set to neutral angles (0 rad). The foot was immobilized on footplate using straps across the instep and metatarsals.
3. Women (quadriceps): Tested was performed with a chair-back angle of 1.48 rad and the knee angle fixed at 1.57 rad to ensure firm tibial contact with the restraint strap.

All individualized joint configurations, limb positions, and dynamometer settings were recorded during a screening session to ensure exact replication across all subsequent visits. In the men, both muscle groups were tested via NMES within the same session. The testing order was randomized between participants but held constant across subsequent visits for each individual. In the women, NMES testing as described below was preceded by determining the torque-velocity relationship during voluntary exercise as reported previously (21).

NMES was performed using a Digitimer DS7AH constant current stimulator (Hertfordshire, UK) to evoke isometric contractions. Testing began by establishing the current required to elicit maximal twitch torque. Single electrical pulses (400 V, 200 μs) were delivered every 3 to 5 s while the current intensity was progressively increased until twitch torque plateaued. Four unpotentiated twitches were then elicited at 1 s intervals, after which the participant performed a 6 s MVIC to induce twitch potentiation. Participants were instructed to attempt to extend their elbow, ankle, or knee as “hard and fast as possible” (23) and strong verbal encouragement was provided during each effort. Four potentiated twitches were then elicited immediately after the MVIC. Participants rested for 10 min rest to allow for reversal of the potentiation and this sequence was repeated.

Following an additional 10 min of rest, the muscle(s) of interest were stimulated for 1 s at 100 Hz and increasing intensity to determine the current needed to elicit approximately 33% of maximal voluntary torque. Trains of stimuli were then delivered at 1, 5, 10, 15, 20, 25, 30, 35, 40, 60, 80, and 100 Hz every 3–5 s to establish the torque-frequency relationship as previously described (14,19).

### Determination of RTD

Torque data were analyzed using Biopac AcqKnowledge version 5.08. Voluntary RTD was determined based on the average slope over the first 0-30 ms, 0-50 ms, and 0-200 ms of the MVICs used to induce twitch potentiation. Contraction onset was manually determined based on the last trough before which torque rose above the background noise of þ0.05 Nm. This approach is considered more reliable than automated methods (24). Maximal voluntary RTD was also calculated based on the 1^st^ derivative of the torque over a 20 ms time interval. The same method was used to determine the maximal tetanic RTD during NMES at 100 Hz. All analyses were performed by a single investigator who was blinded to treatment.

### Statistical analysis

Two-way (i.e., treatment (placebo, NO_3_^−^) x RTD metric (0-30 ms, 0-50 ms, 0-200 ms, maximum)) analyses of variance (ANOVA) were used to determine the effects of dietary NO_3_^−^ on voluntary absolute (i.e., Nm/s) and relative (i.e., as % of maximal isometric torque) RTD in the triceps brachii and triceps surae. Three-way ANOVAs (i.e., treatment x RTD metric x menstrual cycle phase (EF, LF)) were used to compare voluntary absolute and relative RTD in the quadriceps. The effects of treatment on absolute and relative tetanic RTD were determined using paired t-tests and two-way ANOVA (i.e., treatment x menstrual cycle phase) in the triceps brachii/surae and quadriceps, respectively. Statistical analyses were performed using GraphPad Prism 11.1.0 build 224 (GraphPad Software, La Jolla, CA). All data are reported as mean ± standard deviation (SD). Two-tailed P values <0.05 were considered statistically significant. Effect sizes are presented as partial eta squared 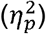.

## RESULTS

### Voluntary RTD

Absolute and relative voluntary RTD in the triceps brachii and triceps surae muscles of the men and the quadriceps muscle of the women are shown in Figs. 1–3, respectively.

**Figure 1.**
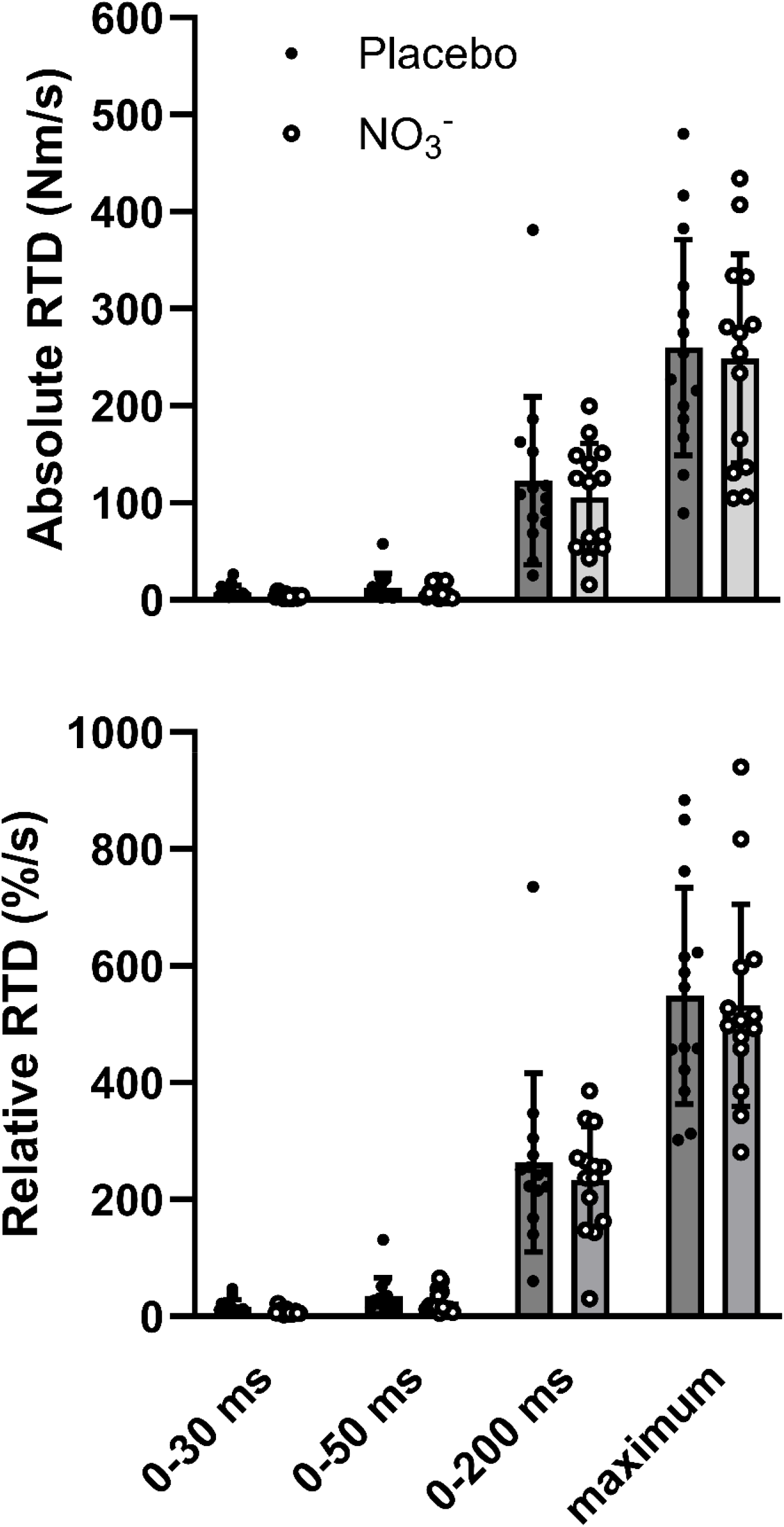
Effect of acute dietary nitrate (NO_3_^−^) supplementation on voluntary absolute (*top panel*) and relative (*bottom panel*) rate of torque development (RTD) in the triceps brachii of n=14 healthy young men.

**Figure 2.**
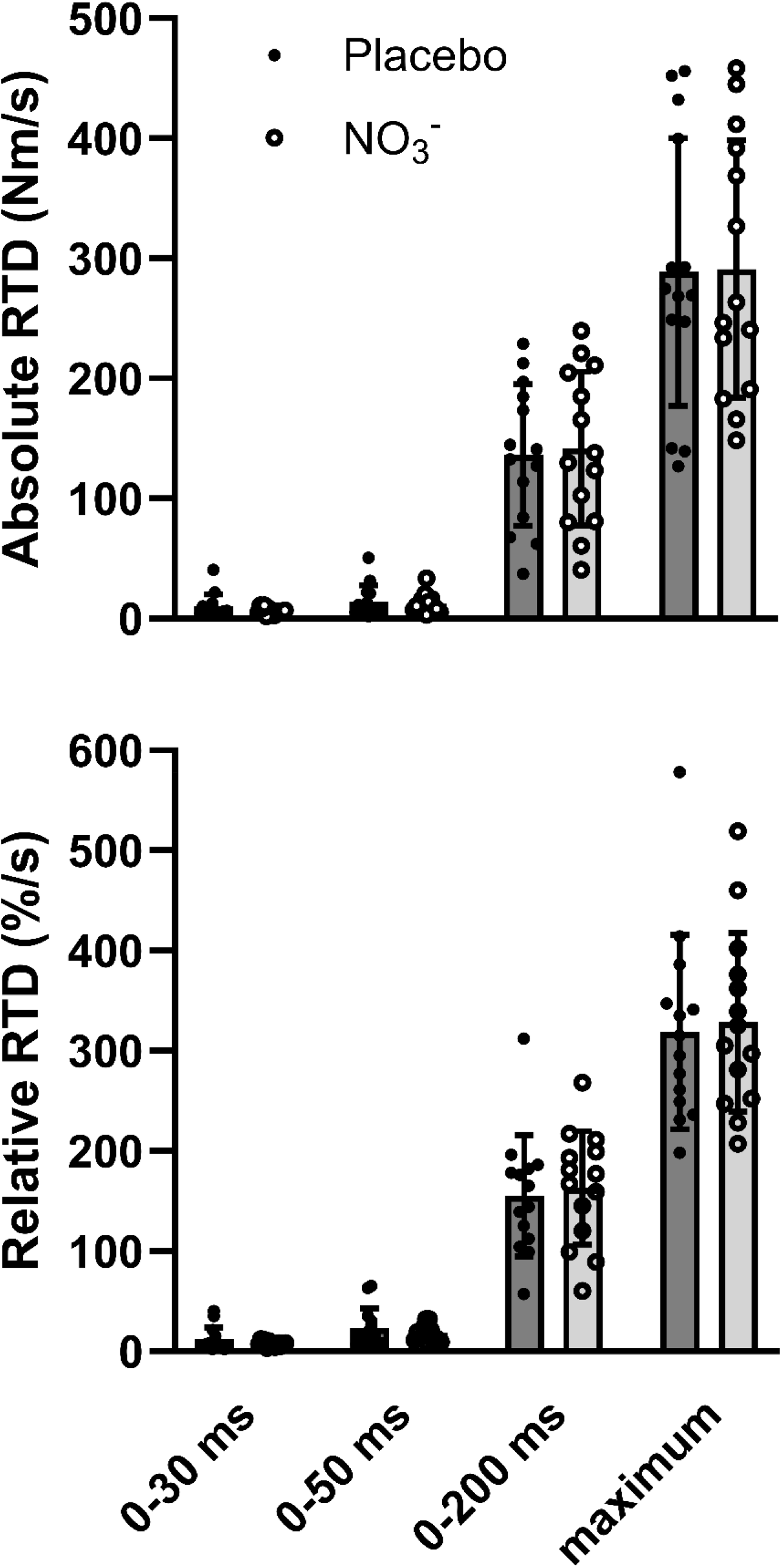
Effect of acute dietary nitrate (NO_3_^−^) supplementation on voluntary absolute (*top panel*) and relative (*bottom panel*) rate of torque development (RTD) in the triceps surae of n=14 healthy young men.

**Figure 3.**
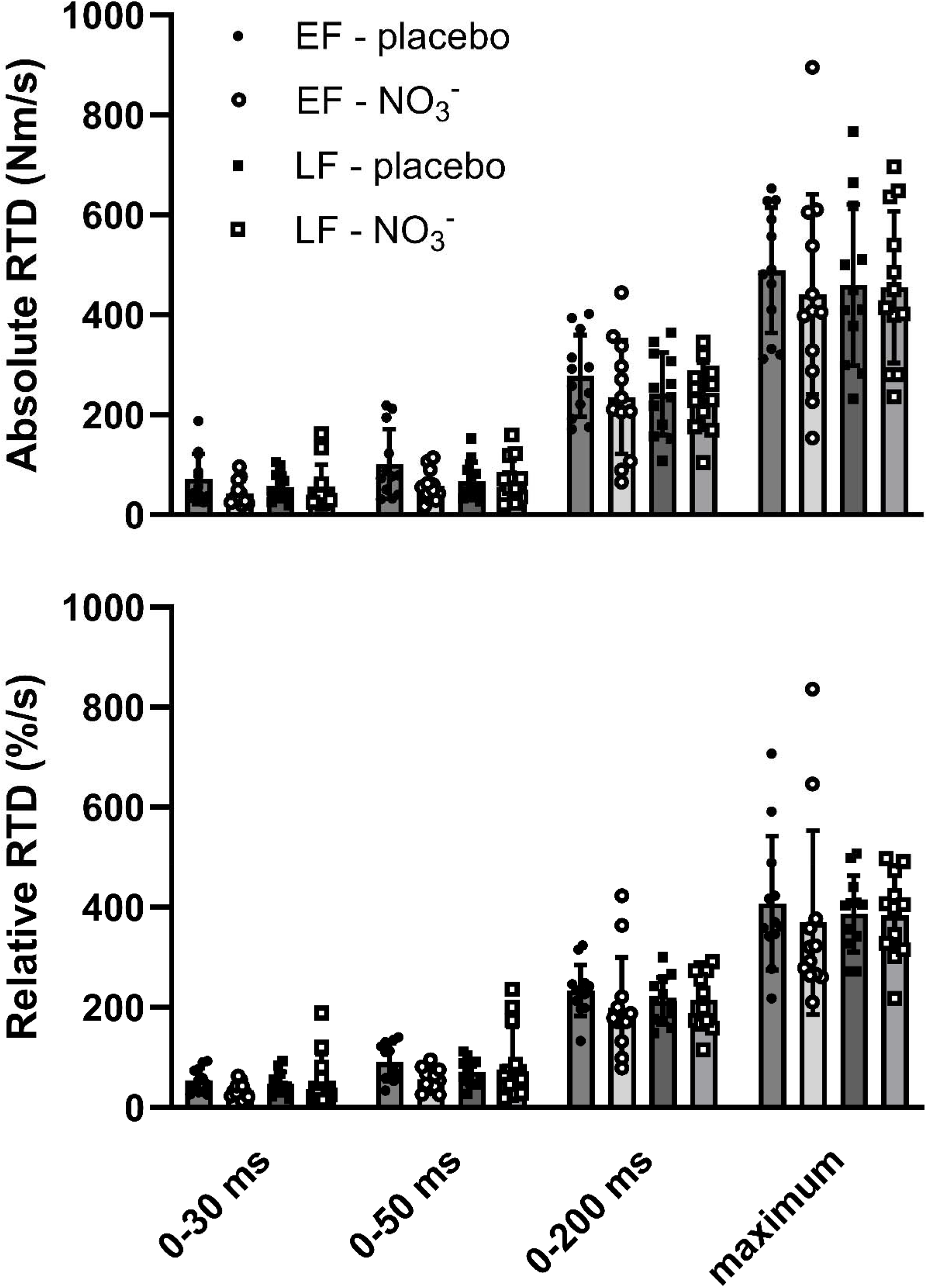
Effect of acute dietary nitrate (NO_3_^−^) supplementation during the early follicular (EF) and late follicular (LF) phases of the menstrual cycle on voluntary absolute (*top panel*) and relative (*bottom panel*) rate of torque development (RTD) in the quadriceps of n=12 healthy young women.

In the triceps brachii, absolute RTD varied significantly based on the metric (P<0.0001, 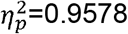, but the treatment and treatment x metric interaction effects were not significant (P=0.3514, 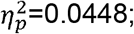 and P=0.7874, 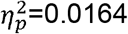, respectively). There was no significant effect of treatment on maximal isometric torque (placebo: 49.6±14.2 Nm, NO_3_^−^: 49.4±15.2 Nm, P=0.9457, 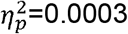). Relative RTD therefore also varied significantly by metric (P<0.0001, 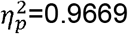), but not by treatment (P=0.3085, 1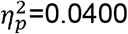). The treatment x metric interaction effect was also not significant (P=0.8099, 1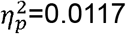).

Similar results were obtained in the triceps surae, with absolute RTD significantly differing based on the metric (P<0.0001, 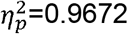), but not treatment (P=0.9639, 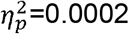) or treatment x metric interaction (P=0.7758, 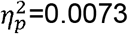). Maximal isometric torque did not differ by treatment (placebo: 98.5±34.2 Nm, NO_3_^−^: 97.4±29.9 Nm, P=0.7995, 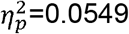). Thus, relative RTD also varied significantly by metric (P<0.0001, 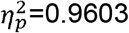), but not treatment (P=0.8875, 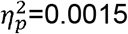) or treatment x metric interaction (P=0.6576, 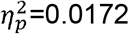).

Finally, in the quadriceps femoris, absolute RTD differed significantly by metric (P<0.0001, 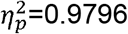), but not by treatment (P=0.5802, 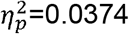) or phase (P=0.1837,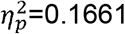). There were also no significant interaction effects (metric x treatment: P=09629, 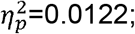; metric x phase: P=0.9629, 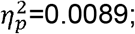 treatment x phase: P=0.1887, 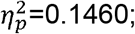 metric x treatment x phase: P=0.9886, 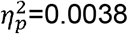). Maximal isometric torque also did not differ by treatment (P=0.1354, 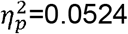), phase (P=0.8086, 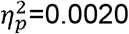), or their interaction (P=0.4476, 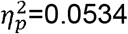). Average values were 128.6±43.4, 124.0±37.0, 119.4±34.4, and 122.5±41.5 Nm for EF-placebo, EF-NO_3_^−^, LF-placebo, and LF-NO_3_^−^ trials, respectively. Consequently, relative RTD also differed significantly by metric (P<0.0001, 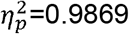), but not treatment (P=0.9262, 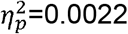), phase (P=0.2688, 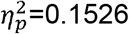), metric x treatment (P=0.9675, 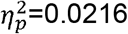), metric x phase (P=0.8318, 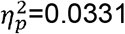), treatment x phase (P=0.0520, 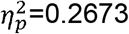), or metric x treatment x phase (P=0.8787, 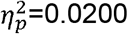).

### Tetanic RTD

Absolute and relative tetanic RTD in the triceps brachii and triceps surae muscles of the men are shown in Table 1. NO_3_^−^ supplementation had no effect on absolute RTD in the triceps brachii (P=0.7522, 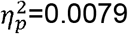) or in the triceps surae (P=0.8930, 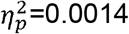). Peak tetanic torque was also the same during both trials in the triceps brachii (placebo: 17.2±5.9 Nm, NO_3_^−^: 17.3±6.0 Nm, P=0.9700, 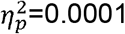) and triceps surae (placebo: 36.2±12.4 Nm, NO_3_^−^: 36.5±9.0 Nm, P=0.8931, 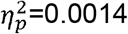). Relative RTD therefore also did not differ in either muscle group (i.e., P=0.8710, 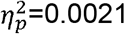 and P=0.8650, 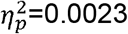, respectively).

**Table 1.** Absolute and relative tetanic RTD in the triceps brachii and triceps surae muscles.

| <u>Muscle and measurement</u> |  | <u>Placebo</u> | <u>NO<sub>3</sub><sup>-</sup></u> |
| --- | --- | --- | --- |
| Triceps<br>brachii | Absolute RTD (Nm/s) | 198.2±69.4 | 193.1±52.7 |
|  | Relative RTD (%/s) | 1207±303 | 1197±354 |
| Triceps surae | Absolute RTD (Nm/s) | 172.5±71.1 | 171.1±53.6 |
|  | Relative RTD (%/s) | 484±120 | 480±122 |

RTD, rate of torque development. Values are mean±S.D. for n=14 healthy young men.

Absolute and relative tetanic RTD in the quadriceps muscle of the women are shown in Fig. 4. Absolute RTD did not differ by treatment (P=0.7039, 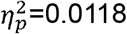), phase (P=0.1255, 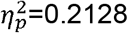), or their interaction (P=0.4989, 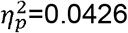). Peak tetanic torque also did not differ by treatment (P=0.0517, 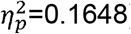), phase (P=0.5646, 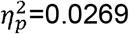), or their interaction (P=0.8342, 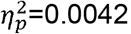), averaging 41.6±17.6, 42.3±11.7, 37.9±14.7, and 39.6±13.8 Nm, in the EF-placebo, EF-NO_3_^−^, LF-placebo, and LF-NO_3_^−^ trials, respectively. As a result, relative RTD also did not differ by treatment (P=0.7093, 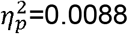), phase (P=0.9572, 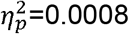), or their interaction (P=0.8640, 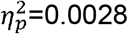).

**Figure 4.**
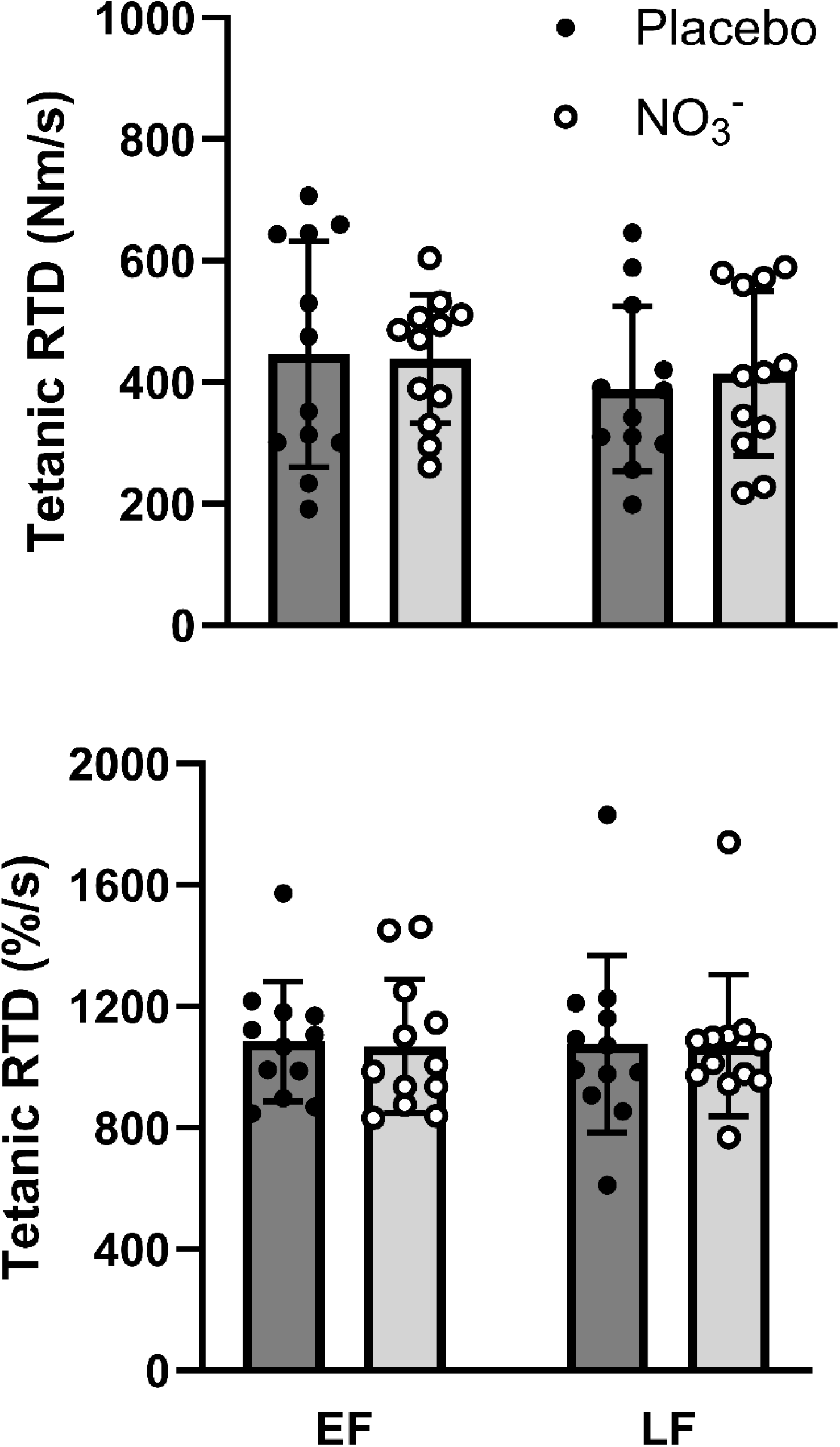
Effect of acute dietary nitrate (NO_3_^−^) supplementation during the early follicular (EF) and late follicular (LF) phases of the menstrual cycle on tetanic absolute (*top panel*) and relative (*bottom panel*) rate of torque development (RTD) in the quadriceps of n=12 healthy young women.

## DISCUSSION

The purpose of the present investigation was to determine the effects of acute NO_3_^−^ supplementation on RTD as influenced by fiber type and sex. We found that dietary NO_3_^−^ had no significant impact on either early- or late-phase/maximal voluntary RTD or on maximal tetanic RTD in either the predominantly fast-twitch triceps brachii or predominantly slow-twitch triceps surae of healthy young men. There was also no influence of NO_3_^−^ intake on these parameters in the quadriceps femoris of healthy young women, in either the EF or LF phase of the menstrual cycle. This is the first study to determine the effects of dietary NO_3_^−^ on RTD in muscles differing in fiber type distribution. It also the first to specifically study young women while controlling for hormonal status.

As previously discussed, some previous studies have found that either acute or repeated NO_3_^−^ supplementation increases voluntary RTD in the mixed quadriceps muscle of younger individuals (18,20,21), whereas others have not (16,17). One possible explanation for these disparate findings is a fiber type specific effect of dietary NO_3_^−^ (14,25), with studies reporting positive results having by chance enrolled participants with a greater percentage of fast-twitch fibers. The present data, however, would seem to rule out this possibility, as NO_3_^−^ intake did not improve voluntary RTD even in the triceps brachii, which is composed of 60-70% fast-twitch fibers (14). This was true during both the early, primarily neurally determined, and later, more muscularly based, phases of voluntary torque development. Importantly, there was also no difference in maximum tetanic RTD, which is independent of central motor drive and hence more reflective of the inherent contractile properties of muscle, especially the crossbridge cycling rate (26). This result is consistent with the findings of Tillin et al. (17), which is apparently the only other study to have determined the effects of NO_3_^−^ intake on this important parameter. We therefore conclude that dietary NO_3_^−^ does not increase either voluntary or tetanic RTD in healthy young men, regardless of muscle fiber type.

Just as there were no differences in the triceps brachii or triceps surae of the healthy young men that we studied, there was also no effect of dietary NO_3_^−^ on voluntary or tetanic RTD in the quadriceps femoris of healthy young women. The present results are therefore consistent with those of Poredoš et al. (18), although unlike the present study these investigators did not control for hormonal status. However, they differ from previous studies that have found that acute (27) or repeated (28) NO_3_^−^ intake increases voluntary RTD in post-menopausal women. This observation again highlights the fact that pre- and post-menopausal women seem to respond differently to dietary NO_3_^−^, as previously discussed (19). Alternatively, however, the contrast between the present results and these previous studies (27,28) may be due to other factors, e.g., the use of blood flow restriction to magnify NO_3_^−^/NO_2_^−^ reduction to NO in the study of Proctor et al. (27). Given the impact of RTD on physical functioning (6,7) and fall risk (8) in older individuals, additional studies of post-menopausal women are warranted to verify the beneficial effect of dietary NO_3_^−^ on RTD in this population. Based on our results and those of Poredoš et al. (18), however, it appears that it does not increase RTD in younger women.

Although dietary NO_3_^−^ has been consistently found to increase maximal muscle speed and power in humans (13), the mechanism by which NO_3_^−^ supplementation enhances human muscle contractility remains to be established. However, the lack of improvement in early phase voluntary RTD in the present and most previous studies (16–18) along with unaltered EMG activity (16) or voluntary activation (19,39-42) argues against changes in neural drive. Likewise, the failure of dietary NO_3_^−^ to increase maximal tetanic RTD (present results; 17) or peak torque during tetanic stimulation (present results; 16,17,36) seemingly rules out changes in the rate of crossbridge cycling or the amount of force generated per crossbridge, respectively. Changes in Ca^2+^ release are another possibility, but both we (14,19) and numerous others (e.g., 16, 37-42) have found that NO_3_^−^ intake has no impact on peak twitch torque, which is inconsistent with such a mechanism. Thus, at present the best explanation for improvements in muscle contractile function due to dietary NO_3_^−^ would seem to be an increase in Ca^2+^ sensitivity, as evidenced by a leftward shift in the torque-frequency relationship (16,36), especially in fast-twitch muscle fibers (14). This could explain why Haider and Folland (16) found that NO_3_^−^ supplementation increased RTD during the first 50 ms of supramaximal nerve stimulation (8 impulses at 300 Hz), while Ca^2+^ levels would have still been increasing (34,35), even though it does not increase maximal RTD during tetanic stimulation (present results; 17), which is reached later, once Ca^2+^ concentrations peak and RTD is limited by the crossbridge cycling rate. If so, however, this increase in Ca^2+^ sensitivity due to NO_3_^−^ intake would have to be due to some mechanism other than phosphorylation of the myosin regulatory light chain (mRLC), as we initially hypothesized (12). This is because we have recently demonstrated that there is no interaction between NO_3_^−^ supplementation and post-activation twitch potentiation (14), which is also largely due to mRLC phosphorylation (30,31).

Accurate measurement of voluntary RTD can be challenging due to high between-trial variability (2). Thus, a limitation of the present investigation is that our conclusions are based upon only two replicate measurements per condition (i.e., placebo vs. NO_3_^−^), albeit in two different muscle groups in men (14) and two different phases of the menstrual cycle in women (19). In contrast, others have assessed the effects of NO_3_^−^ on voluntary RTD using 3-15 “explosive” contractions (16–18). However, except for the increase in maximal voluntary RTD observed in men only by Poredoš et al. (18), our results are consistent with these previous studies. We also found no changes in either early phase (data not shown) or maximal tetanic RTD, thus reinforcing our findings during voluntary exercise. Nonetheless, it is possible that we may have overlooked a small effect of dietary NO_3_^−^ on either early phase or maximal voluntary RTD. Other limitations of the present study include that we only determined the effects of a single, acute dose of NO_3_^−^, and did not test varying doses (20) or the effects of repeated supplementation (21). Our conclusions are also limited to the effects of dietary NO_3_^−^ on RTD in healthy young men and women, since as previously discussed older individuals may respond differently.

## CONCLUSION

In summary, we have determined the effects of acute NO_3_^−^ supplementation on early phase and maximal voluntary and maximal tetanic RTD in the triceps brachii and triceps surae of men and the quadriceps femoris of women, the latter during both the EF and LF phases of the menstrual cycle. There were no changes in any parameter under any condition. These data argue against changes in neural activation or crossbridge cycling rate as potential mechanisms by which dietary NO_3_^−^ may enhance the contractility of human muscle.

## FUNDING

Madison J. Fry was supported by an Undergraduate Research Opportunity Program award from the Center for Research and Learning at Indiana University Indianapolis during performance of the primary investigation of young healthy women. She is presently affiliated with the Department of Nutrition and Exercise Physiology at the University of Missouri-Columbia, Columbia, MO 65211-7703. This secondary analysis was supported by grant R33 HL155858 to Drs. Linda R. Peterson, Andrew R. Coggan, and Kenneth B. Schechtman from the National Heart, Lung, and Blood Institute of the National Institutes of Health.

## DISCLOSURE STATEMENT

No conflicts of interest, financial or otherwise, are declared by the authors.

## DATA AVAILABILITY STATEMENT

Data are available from the corresponding author upon reasonable request.

## Notes

### Competing Interest Statement

The authors have declared no competing interest.

